# Altered lipid homeostasis underlies selective neurodegeneration in SNX14 deficiency

**DOI:** 10.1101/2022.11.30.516463

**Authors:** Yijing Zhou, Vanessa B. Sanchez, Peining Xu, Marco Flores-Mendez, Brianna Ciesielski, Donna Yoo, Hiab Teshome, Mike Henne, Tim O’Brien, Clementina Mesaros, Naiara Akizu

## Abstract

Dysregulated lipid homeostasis is emerging as a potential cause of neurodegenerative disorders. However, evidence of errors in lipid homeostasis as a pathogenic mechanism of neurodegeneration remains limited. Here, we show that the cerebellar neurodegeneration caused by SNX14 deficiency is associated with lipid metabolism defects. Recent *in vitro* and *in silico* studies indicate that SNX14 is an inter-organelle lipid transfer protein that regulates lipid droplet biogenesis and fatty acid desaturation, suggesting that human SNX14 deficiency belongs to an expanding class of cerebellar neurodegenerative disorders caused by altered cellular lipid homeostasis. To test this hypothesis, we generated a mouse model that recapitulates the human SNX14 deficiency at genetic and phenotypic level. Through histological and transcriptomic analyses, we demonstrate that cerebellar Purkinje cells are selectively vulnerable to SNX14 deficiency, while forebrain regions preserve their neuronal content. Ultrastructure and lipidomic studies reveal widespread lipid storage and metabolism defects in SNX14 deficient mice. Furthermore, we identify a unique lipid metabolite profile that links the accumulation of acylcarnitines with the selective cerebellar neurodegeneration in SNX14 deficiency. These findings highlight the importance of lipid homeostasis for neuronal function and survival and suggest a mechanism for selective cerebellar vulnerability to altered lipid homeostasis.

## Introduction

Neurodegenerative disorders are characterized by a progressive loss of selective neuronal types often associated with the accumulation of toxic protein aggregates (*1*). To better understand disease mechanisms and find therapeutic alternatives, the field has principally focused on the study of protein quality control pathways, including autophagy (*2, 3*). In contrast, little attention has been paid to lipid homeostasis pathways despite their well-stablished association with neurodegeneration and evidence showing their relevance for the integrity and function of cellular organelles (*4–7*).

Genetic disorders affecting regulators of lipid homeostasis often show neurodegeneration particularly affecting the cerebellum and spinal cord (*8, 9*). The cerebellum integrates motor function with cognition, emotion and language, and its dysfunction is documented in a wide spectrum of neurological disorders (*10–12*). Among cerebellar disorders, childhood onset spinocerebellar ataxias are the most severe. In addition to impaired motor coordination and balance, spinocerebellar ataxia in children is often accompanied by additional neurologic and systemic symptoms, including neurodevelopmental delay and intellectual disability (*13, 14*). Recent efforts that combine patient registry assemblies with advances in sequencing technologies are revealing a new class of childhood cerebellar neurodegenerative disorders caused by disfunction of lipid homeostasis pathways (*8, 15*).

Mutations in *Sorting Nexin 14 (SNX14)* are the cause of a childhood-onset ataxia known as Spinocerebellar Ataxia Recessive 20 (SCAR20), characterized by a progressive cerebellar degeneration and severe intellectual disability (*16–18*). We previously discovered that SCAR20 is associated with enlarged lysosomes and altered autophagy in neural cells derived from patients (*16*). These findings were also reproduced in patient skin fibroblasts and SNX14 deficient U2OS cell lines but deemed secondary to defects in cholesterol distribution and neutral lipid metabolism (*19*). Subsequent studies identified SNX14 as a regulator of cholesterol homeostasis in two independent genome wide perturbation screens (*20, 21*). Although the mechanisms by which SNX14 regulates cholesterol trafficking is still unknown, recent reports demonstrate that SNX14 is recruited to the endoplasmic reticulum (ER)-lipid droplet (LD) contact sites to facilitate the incorporation of fatty acids (FA) into triglycerides of growing LDs (*22*). In this process, SNX14 interacts with SCD1, an ER anchored fatty acid desaturase, to cooperate in FA desaturation and incorporation into LDs (*23*). Consequently, SNX14 deficient cells show enhanced toxicity to saturated fatty acids and defective FA-stimulated LD biogenesis (*22, 23*). Furthermore recent structural analyses facilitated by AlphaFold2 predictions suggest that SNX14 and its SNX-RGS family members may be involved in intracellular lipid transfer (*24*). However, it is currently unknown if the role of SNX14 in lipid homeostasis regulation is implicated in the pathogenesis of SCAR20.

To shed light on the cellular and molecular mechanisms that lead to cerebellar degeneration and intellectual disability in SNX14 deficiency, we generated the first *Snx14* full body knock out mouse (*Snx14* KO) that survives to adulthood. Our work shows that *Snx14* KO mice recapitulate cerebellar atrophy, and motor and cognitive defects of SCAR20 patients. Whereas cerebellar atrophy is associated with Purkinje cell (PC) degeneration, forebrain regions responsible for cognitive behavior remain protected from neurodegeneration. Guided by transcriptomic analyses that pointed to lipid dysregulation as a potential cause of selective cerebellar degeneration, we identify tissue specific alterations of lipid profiles in *Snx14* KO mice tissue. Particularly, *Snx14* KO cerebral cortices exhibit reduced phosphatidylethanolamines (PEs) levels that may be associated with synaptic dysfunction, while accumulation of Acylcarnitines (AcCa-s) is unique to pre-degenerating cerebella and likely associated with selective cerebellar neurodegeneration. Finally, we show that SNX14 deficiency reduces lipid droplet (LD) numbers in the liver and causes lipid storage defects in cerebellar PCs. Overall, our work provides evidence for the involvement of lipid homeostasis defects in selective neurodegeneration and uncovers lipid targets for therapeutic interventions.

## Results

### SNX14 deficiency causes partial embryonic lethality and developmental delay in mice

SCAR20 patients share clinical features of developmental delay and perinatal onset neurodegeneration of the cerebellum. Previous work suggested that the severity of the developmental phenotypes is species-specific, with SNX14 deficient mice showing fully penetrant embryonic lethality, while dogs and zebrafishes displaying neurological and metabolic defects reminiscent of human SCAR20 patients (*25*). However, by randomly introducing a frameshift 1 bp deletion in the exon 14 of *Snx14* (c.1432delG; p.Glu478Argfs*18) we successfully generated SNX14 deficient mice (*Snx14* KO) that are viable and thrive despite a complete loss of SNX14 protein and 90% reduction of the transcript when the mutation is in homozygosity (Fig. 1A and Fig S1A-D).

**Fig. 1.**
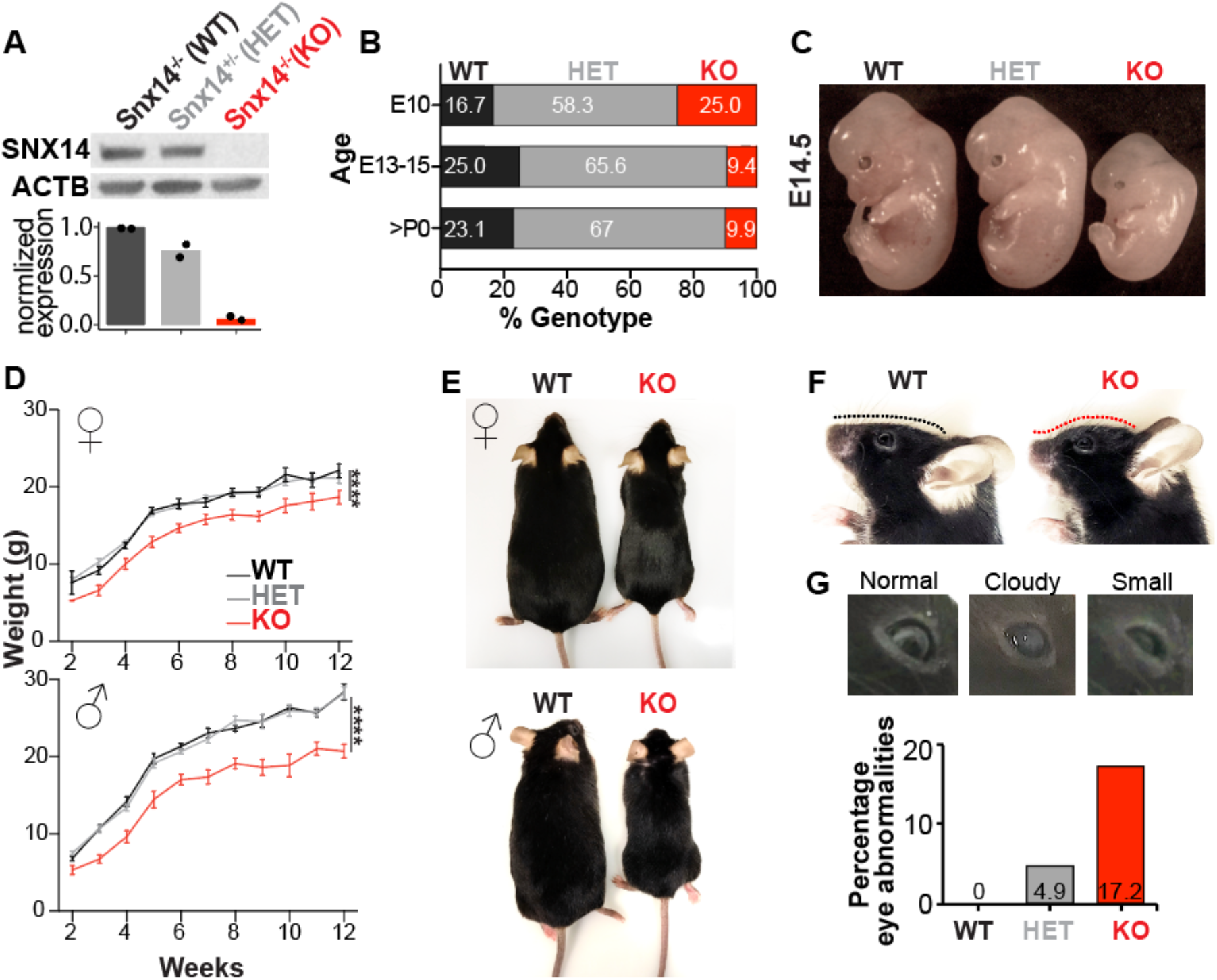
SNX14 deficient mice show developmental delay and atypical facial features. (**A**) Representative western blot (WB) analysis showing loss of SNX14 expression in *Snx14* KO mice tissue. Beta-actin is used as loading control. Bar graphs show WB band densitometry quantification of SNX14 relative to ACTB. N=2. (**B**) Percentage of embryos/mice with the indicated genotypes obtained from heterozygous parent matings, Chi-square test shows significant discrepancy between >P0 observed and expected values (P value=0.001) indicating embryonic lethality of KOs. E10, N=12; E13-15, N=73; >P0, N=91. (**C**) Representative image of WT, HET, and KO E14.5 embryos showing smaller size of KOs. (**D**) Growth curve showing consistently lower body weight of 2-12 week-old *Snx14* KO males and females. Graph shows average ±S.E.M of N≥3. Two-way ANOVA test shows significant effect of genotype (****P<0.0001). (**E**) Representative images of 9-month-old WT and KO littermates of each gender. (**F**) Representative images showing the atypical face with forehead protrusion of KO mice (red line) compared to a WT littermate. Mice shown were 6 mo. (**G**) Representative images showing eye abnormalities, including cataracts (cloudy), and microphthalmia (small) of KO mice. Mouse shown were 8 mo. Bar graph shows percentages of mice with eye abnormality for each genotype.

Although *Snx14* KO mice survived to adulthood, we noticed that they were born in a lower than the expected Mendelian ratio (observed 9.9% vs expected 25%) (Fig 1B). To test if the reduced birth ratio was due to embryonic lethality, we genotyped embryos produced by heterozygous breeding pairs and uncovered that about half of *Snx14* KO embryos die between embryonic day (E)10 and E15. The other half were easily distinguishable by their smaller size, a feature that persisted throughout neonate and adulthood period (Fig 1C-E). Notably, similar to SCAR20 patients (*16–18*), adult *Snx14* KO mice showed dysmorphic facial features characterized by upturned nose, bulging forehead and eye defects (Fig 1F, G). These data indicate that SNX14 deficiency in mice causes developmental delay phenotypes reminiscent of SCAR20.

### *Snx14* KO mice have motor and cognitive behavioral defects

Unlike SCAR20 patients who show severe gait abnormalities typical of cerebellar degeneration, *Snx14* KO mice were undistinguishable from their wild type (WT) littermates based on their homecage walking activity. However, Catwalk gait analysis revealed a mild gait disruption characterized by longer paw stand time and faster swing speed of the limbs (Fig 2A, B). Further, functional gait disruption was seen on the horizontal Metz ladder, where mice cross a series of rungs separated by varying distances. Here, *Snx14* KO mice had significantly more foot slips than control mice (Fig 2C). Moreover, *Snx14* KO mice underperformed when challenged with complex motor tasks that require coordination and balance. On the accelerating Rotarod, *Snx14* KO mice showed difficulty maintaining balance (Fig 2D), similar to other cerebellar ataxia mouse models (*26*). In addition, the accelerating Rotarod procedure was performed in three consecutive days to assess motor learning. Remarkably, while WT mice improved their performance over trial, *Snx14* KO learning rate was low, especially for females (Fig 2E).

**Fig. 2.**
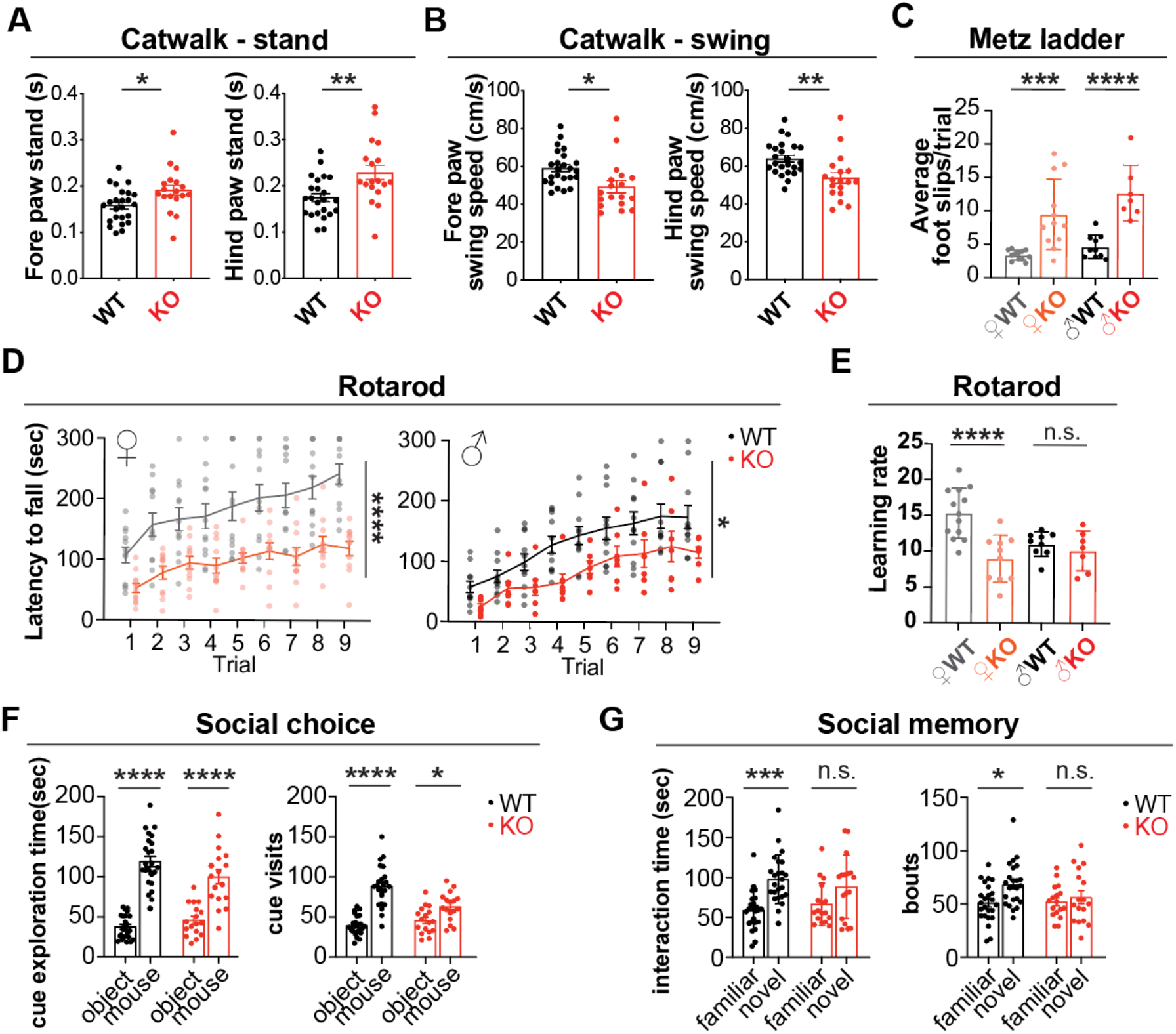
SNX14 deficiency in mice recapitulates motor and behavioral deficits of SCAR20. **(A-B)** Catwalk analysis showing altered gait of KO mice evidenced by comparison of stand (A) and swing (B) with WT mice. Bar graph shows average ±S.E.M of n=24 WT and n=18 KO mice. Welch’s *t*-test (*P<0.05, **P<0.01). (**C**) Metz ladder rung test showing altered limb placing and coordination of KO males and females. Bar graph shows average foot slip of 5 trials performed in consecutive days ±S.E.M of n=10 WT males, n=7 KO males, n=12 WT females, n=12 KO females. Two-way ANOVA followed by Sidak’s post hoc test (***P<0.001, ****P<0.0001). (**D**) Accelerating rotarod test revealing defects in motor performance of KO mice in the 9 trials performed over 3 consecutive days. Graph shows average latency to fall ±S.E.M of n=11 WT males, n=7 KO males, n=13 WT females, n=11 KO. Two-way ANOVA test shows significant effect of genotype (*P<0.05, ****P<0.0001) (**E**) KO females show impaired learning rate on accelerating rotarod performance over time (between trial 1 and 9). Graph shows average learning rate ±S.E.M of n=9 WT males, n=7 KO males, n=13 WT females, n=10. Two-way ANOVA test followed by Sidak’s post hoc test (****P<0.0001, n.s.=non significant). (**F**-**G**) Three-chamber social interaction test showing similar preference for a mouse over an object between WT and KO mice (F) but impaired social novelty preference in KO mice (G). Bar graphs show average and S.E.M of WT n=24, KO n=17. Two-way ANOVA followed by Tukey’s multiple comparisons test ((*P<0.05, ***P<0.001, ****P<0.0001).

Given that intellectual disability accompanies the cerebellar ataxia as hallmark of SCAR20, we wondered whether *Snx14* KO mice had broader behavioral deficits. To answer this question, we performed a test for social preference and recall (*27*). During a choice phase of the procedure, the *Snx14* KO mice showed typical preference for a social cue relative to an inanimate object. Remarkably, during the recall phase the *Snx14* KO mice failed to discriminate between a familiar and a stranger mouse (Fig 2G). Thus, *Snx14* KO mice showed similar preference to the social cue, but their lack of preference toward exploration of the novel mouse suggests a social memory deficit likely caused by dysfunction of brain regions, including the cerebellum (*28, 29*).

### Behavioral defects are associated with cerebellar atrophy

Having established that SNX14 deficient mice recapitulate developmental, motor and behavioral delays of SCAR20, we looked for the underlying neuropathologic causes. In mice, like in humans, SNX14 is widely expressed in the developing and adult brain, with a slight enrichment in older brains (Fig S2A). In line with the expression pattern, the gross brain morphology of *Snx14* KO mice appeared normal during the first month of life but showed defects as mice became older. Specifically, we found that *Snx14* KO mice had smaller cerebellum than WT littermates starting at 2.5 months of age (Fig 3A-B). Interestingly, we did not find remarkable differences in the gross morphology or size of other brain areas (Fig 2C), suggesting that the cerebellum is particularly vulnerable to SNX14 deficiency.

**Fig. 3.**
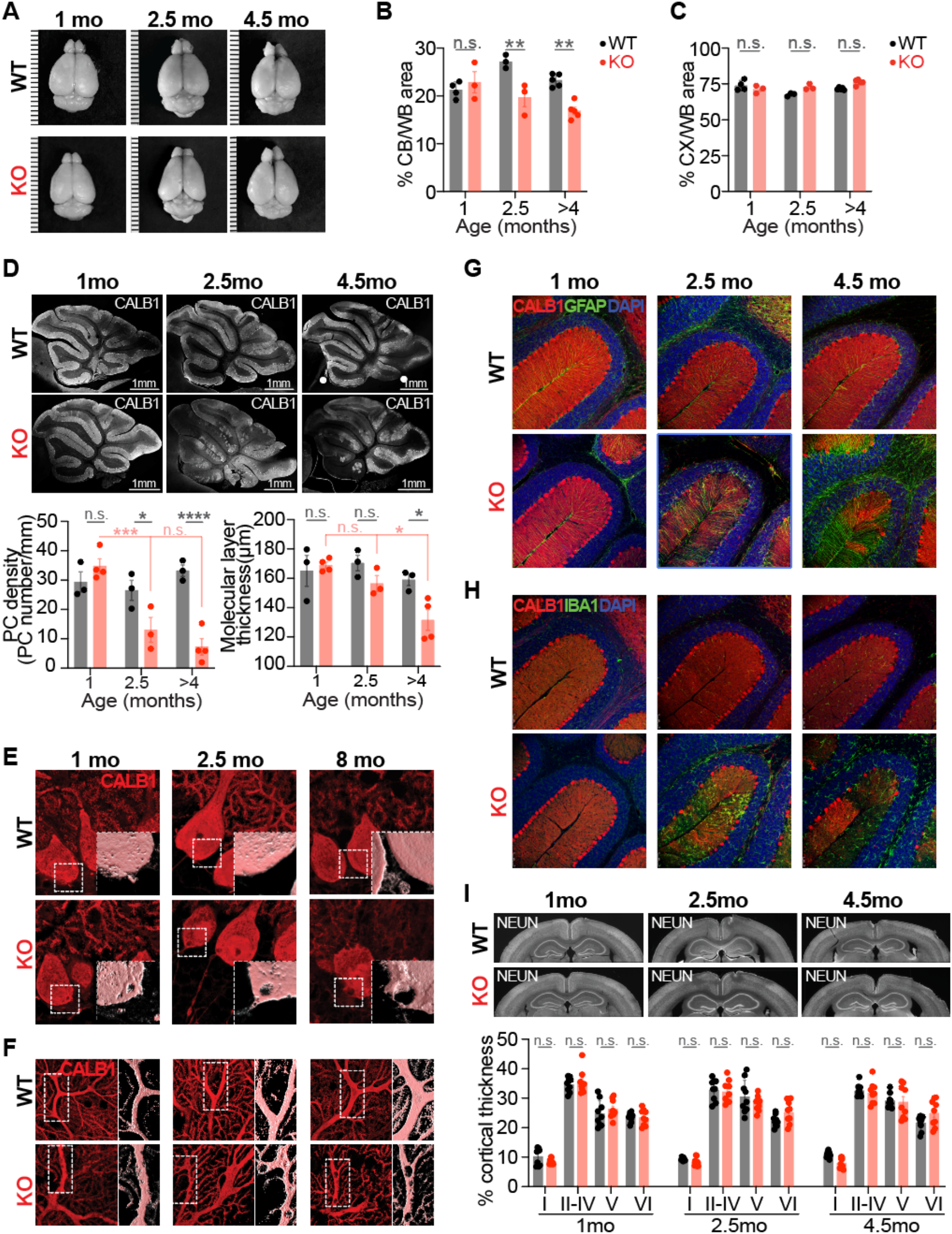
SNX14 deficiency causes selective cerebellar degeneration. (**A**) Representative images of brain from mice with indicated age and genotype. Ruler marks separated by 1mm. (**B-C**) Quantitative comparison of percentage area of cerebellum (CB) (B) or cortex (CX) (C) relative to the whole brain (WB) shows a selective shrinkage of KO CB over time. Bar graphs show average ±S.E.M of N=3-5 mice. Two-way ANOVA followed by Sidak’s test (**P<0.01, n.s.=non significant). (**D**) Cerebellar sagittal sections immunostained with PC specific CALB1 antibody reveal progressive loss of PCs in KO mice. Graphs show average linear density of PCs (right) and average thickness of the molecular layer (left) of Cerebellar Lobule III in n=3-4 mice per genotype and timepoint. Two-way ANOVA followed by Sidak’s test (*P<0.05, ****P<0.0001, n.s.=non significant). (**E-F**) Immunofluorescent staining of PCs with CALB1 antibody reveals progressive accumulation of vacuoles in KO mice PC soma (E) and dendrites (F). (**G-H**) Immunofluorescent staining of astrocytes with anti-GFAP (G) and microglia with anti-IBA1 (H) in green show a progressive accumulation of astrocytes and microglia in degenerating KO cerebella. CALB1 (red) and DAPI (blue) are also shown. The base of Lobule III & IV is shown as representative image. (**I**) Coronal sections of cerebral cortices immunostained with anti-NeuN do not show differences between WT and KO mice. Bar graphs show average ±S.E.M of the percentage thickness occupied by each of the cortical layers (I-VI) in 4-5 cortical regions of 2 mice per genotype and age. Two-way ANOVA followed by Sidak’s test (n.s.=non significant).

### SNX14 deficiency causes selective PCs degeneration

To further determine vulnerabilities of SNX14 deficiency at cellular level, we analyzed cerebellar and forebrain tissue histologically. Within the cerebellum, *Snx14* expression is enriched in Golgi and Purkinje Cells (PCs) (*30*) (Fig S2B). The PCs are some of the largest neurons in the nervous system and their loss is a hallmark of cerebellar ataxias (*31*). Thus, we first analyzed PCs in 1, 2.5 and 4 months old cerebellar sections by immunostaining with Calbindin 1 (CALB1) antibody. At 1 month of age, both WT and *Snx14* KO cerebellar stainings showed perfectly aligned somas in the PC layer and PC dendrites extended into the molecular layer. However, by 2.5 months of age, patches of missing PCs were evident in *Snx14* KO cerebella (Fig 3D). Quantification of PC number per mm of PC layer confirmed significantly lower PC density in lobule III of 2.5 and 4 months old *Snx14* KO cerebella compared to WT (Fig 3D left graph). The loss of PCs in *Snx14* KO cerebella was followed by a reduced thickness of the molecular layer, which was first detectable at 4 months of age (Fig 3D right graph). A closer look to the CALB1 staining, revealed the presence of vacuole-like structures within *Snx14* KO PC dendrites and soma (Fig 3E, F). Although, these vacuoles were more abundant and larger in older cerebella, they were sparsely detected them in 1 month old PCs, suggesting that the vacuoles may be a pathological sign that precedes PC neurodegeneration.

Given that PC degeneration is often followed by disorganization of Bergmann Glia (BG) processes and astrogliosis, we also immunostained sagittal cerebellar sections with anti-GFAP and anti-IBA1 antibodies. Concurrent with the PC loss, anterior lobes of 2.5 month old *Snx14* KO cerebella showed abnormal branching of GFAP positive BG processes (Fig 3G) and an accumulation of IBA1 positive microglia within the molecular layer (Fig 3H). Moreover, we found that reactive astrocytes progressively accumulate nearby the PC layer from 2.5 to 4.5 months of age (Fig 3G). Interestingly, these findings were specific of the anterior lobes of the *Snx14* KO cerebella, while posterior lobes (VIII and IX) did not show signs of neurodegeneration until 11 months of age (Fig S2D, E). Similarly, we never saw loss of neurons or signs of astrogliosis in cortical and hippocampal regions of the forebrain (Fig 3I and Fig S3A-C).

Taken together, our results indicate that despite the wide expression of SNX14 in the whole brain, the forebrain and posterior cerebellum are protected from neurodegeneration, while anterior PCs are selectively affected by SNX14 deficiency, which cause their death after 2 month of age.

### Lipid response genes are dysregulated in pre-degenerating *Snx14* KO mice cerebella

To gain insights into the molecular mechanisms of selective cerebellar PC degeneration, we next analyzed the transcriptome of *Snx14* KO mice cerebella at pre and post-degenerating stages (<2 month old and 1 year old respectively) and compared them with cerebral cortices, which do not show signs of neurodegeneration in our histological studies. After RNA sequencing we defined differentially expressed genes (DEG) those showing absolute log_2_(FC)>0.58 with p-adj<0.05 between *Snx14 KO* and WT tissue. As expected, *Snx14* was downregulated in all *Snx14 KO* samples (Fig 4A, C and Fig S1D). Other than this, differences between *Snx14 KO* and WT cerebral cortex transcriptomes were minor at <2 month of age (4DEGs including *Snx14*) and showed only 38 downregulated and 4 upregulated DEGs at 1 year of age (Fig 4A). None of these DEGs suggested changes in specific cell type composition, which is consistent with the lack of neurodegeneration or neuroinflammation in our histological analyses. We then tested if cortical DEGs were enriched in specific cellular and molecular functional annotations. Given the short list of DEGs at <2 month old cortices, we only performed functional annotation analysis on the 38 downregulated genes at 1 year of age. Results revealed a significant enrichment for genes involved in synaptic function (i.e. synaptic signaling, neurotransmitter transport, GPCR signaling) (Fig 4B). In line with this result, previous work showed that SNX14 promotes synaptic activity in mouse cortical neuronal cultures (*32*). Therefore, although *Snx14* KO cortical neurons do not degenerate, SNX14 deficiency likely affects synaptic activity of forebrain cortical neurons.

**Fig. 4.**
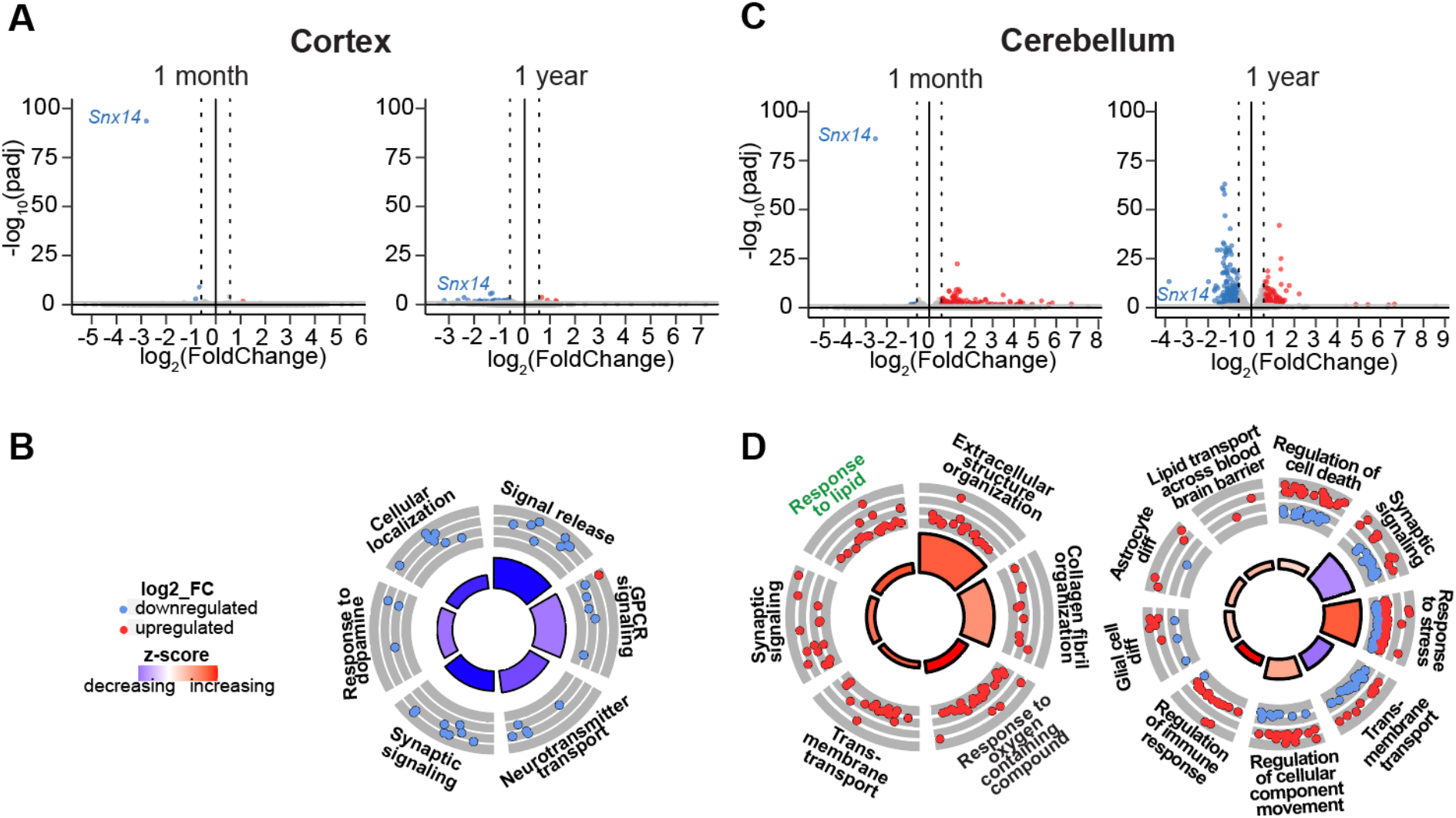
Genes involved in lipid response are differentially expressed in SNX14 deficient cerebella. (**A**) Volcano plots for the analysis of differentially expressed genes (DEGs) showing few DEGs in 1-month and 1-year old *Snx14* KO vs WT cerebral cortices. Dotted vertical lines mark log_2_FC=0.58 and −0.58. Red dots label upregulated DEGs and blue dots downregulated DEGs. (**B**) Circular plot showing selected biological processes (BP) enriched in DEGs of 1-year KO cortices. The outer circle is a scatter plot showing the log_2_(FC) of DEGs in each category (red=upregulated, blue=downregulated). The inner circle is a bar-plot, with the height of the bar indicating the significance of the term (-log_10_(p-adj)) and the color displays the z-score, which represents the average expression level of all DEGs in each category. (**C**) Volcano plots showing DEGs of 1-month and 1-year *Snx14* KO vs WT cerebella. (**D**) Circular plot showing selected BPs enriched in DEGs of 1-month and 1-year old KO cerebella.

Unlike cerebral cortices, cerebellar transcriptomes were markedly different between *Snx14* KO and WT mice, with 158 upregulated and 6 downregulated DEGs at 1 month of age and 169 up- and 226 downregulated DEGs at 1 year *Snx14* KO cerebella. We reasoned that the increase in the amount of downregulated DEGs from 1 month to 1 year of age could reflect the progressive PC loss in *Snx14* KO cerebella. Accordingly, some of the most downregulated DEGs in 1 year *Snx14* KO cerebella correspond to well-known PC markers, such as *Calb1*, *Pcp2, Car8* and *Rgs8*. To unbiasedly test this observation, we compared 1 year old DEG list with a recently reported mouse cerebellar single nuclear RNAseq dataset (*30*). Results confirmed that downregulated DEGs are enriched in genes predominantly expressed in PCs (Fig S4). On the other hand, most of the upregulated DEGs are genes sparsely expressed across various cerebellar cell types, with a group of them typically expressed in astrocytes and macrophage/microglia of the cerebellum respectively (Fig S5). Notably, 1 month old DEGs did not overlap with PC or astroglia specific markers indicating a later onset of neurodegeneration, which agrees with our histological analyses.

Given the lack of neurodegenerative signs in histology or transcriptomic data (Fig 3 and 4), we anticipated that DEGs at 1 month old cerebella could point us to the molecular causes that precede PC neurodegeneration. Like for the cerebral cortex datasets, functional annotation analyses showed an enrichment in synaptic signaling genes (Fig 4D). In addition, the enrichment in “Response to lipids” functional group called our attention because it would suggest that lipid homeostasis defects may precede selective cerebellar neurodegeneration.

### SNX14 deletion alters lipid metabolite levels in a tissue specific manner

We next set out to analyze lipid metabolite composition of pre-degenerating cerebella in <2 month old *Snx14* KO mice and their WT littermates by unbiased lipidomic analysis. As a control of a non-degenerating tissue, we also included their cerebral cortices in the analysis. In addition, since the liver is a lipid rich organ with high content of triglycerides (TGs) stored in lipid droplets, we included liver lipid extracts as a control for lipid metabolite detection. Finally, to distinguish tissue specific lipids from those circulated by their blood supply, we also extracted plasma lipids from circulating blood.

The lipid extracts were analyzed by ultraperformance liquid chromatography-high resolution mass spectrometry (UPLC-HRMS) as previously described (*33*) and after normalization with lipid internal standards, we quantitatively identified more than 200 lipid species per sample. Overall, *Snx14* KO and corresponding WT tissues had similar total lipid concentration (Fig S5A). Moreover, each of the four tissues we analyzed were distinguishable by their relative lipid class abundance. For instance, liver displayed the highest abundance of triglycerides (TG) while the cerebral cortex and cerebellum, both with similar lipid content, had phosphatidylcholines (PCh) as the most abundant lipid class (Fig S5B). This data is consistent with data in the literature (*34, 35*), thus validating our methodology.

Next, we aimed to determine how SNX14 deficiency affects tissue specific lipid composition. To this end, we compared the concentration of each lipid specie in *Snx14* KO and the corresponding WT tissue. Given the role of SNX14 facilitating the incorporation of fatty acids (FA) into triglycerides (TG) during LD biogenesis (*22*), we hypothesized that SNX14 deficiency would result in a depletion of TG levels. Although TGs were undetectable in all cerebellar and cortical samples, *Snx14* KO livers displayed a significant reduction of TGs (Fig 5A and FigS5B), further confirming our hypothesis and reliability of our lipidomic analysis.

**Fig 5.**
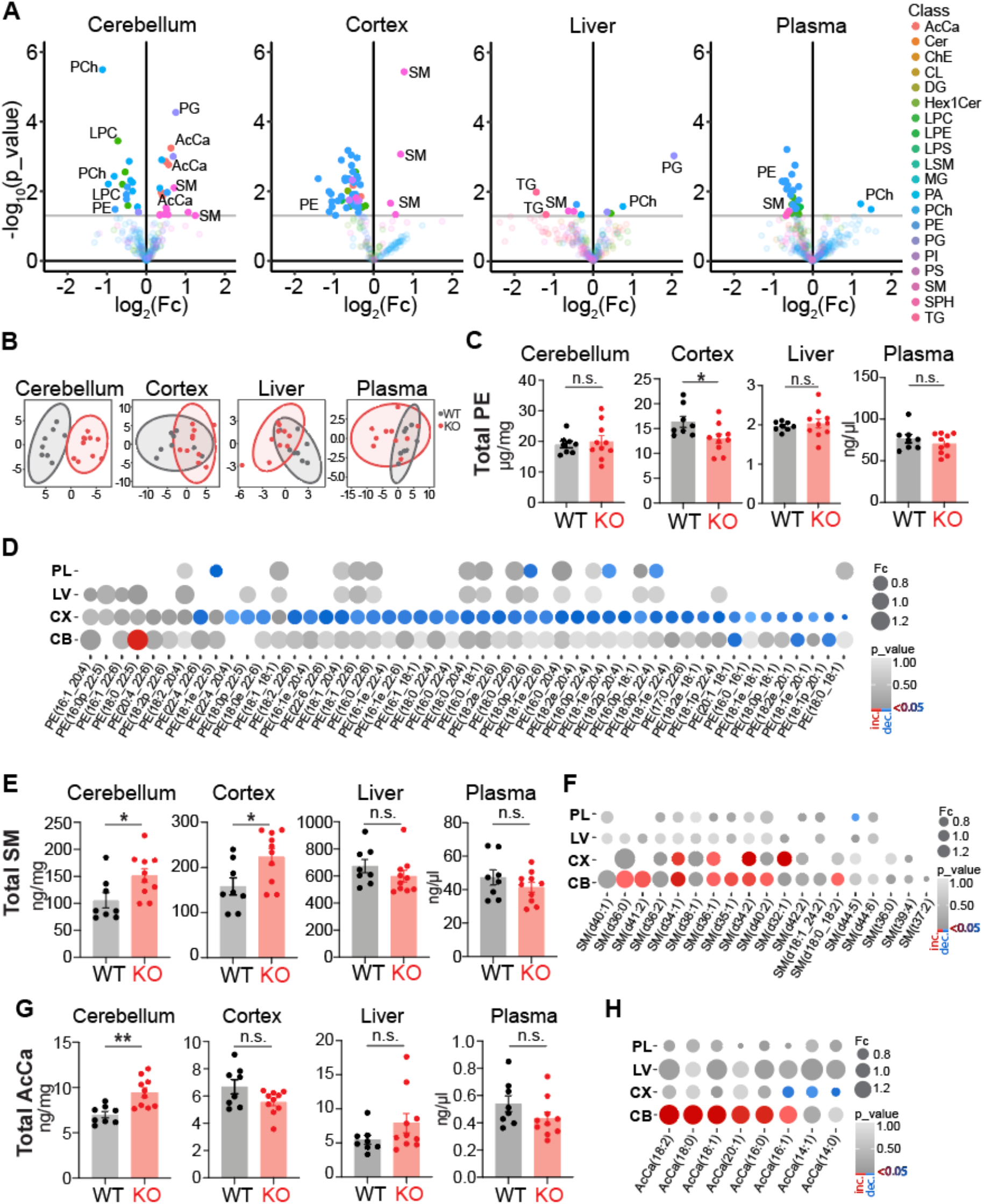
Unique deregulation of lipid metabolites in pre-degenerating KO cerebella. (**A**) Volcano plots showing significantly deregulated lipids in *Snx14* KO cerebellum, cortex, liver, and plasma of 1-2 month old mice, with Pval<0.05 a cut-off (horizontal grey line). Data shows increased concentrations of Acylcarnitine (AcCa) species and specifically in the KO cerebellum. Significantly deregulated key lipid species are labeled. (**B**) Principal component analysis of deregulated lipid species, separates WT and KO samples in two clusters only in the cerebum. **(C)** Bar graphs showing the average ±S.E.M of the sum of PE species concentrations per tissue in n=8 WT and n=10 KO mice. Student *t*-test. (*P<0.05, n.s.=non significant). PEs are significantly reduced in KO cerebral cortices (CX). (**D**) Dotplot depicting fold change (FC) (proportional to dot size) and p-value (in grey intensity scale) of PE species detected in cerebral cortices for all analyzed tissues. Red dots represent significantly increased lipids and blue dots significantly decreased. (E) Bar graphs showing the average ±S.E.M of the sum of SM species concentrations per tissue in n=8 WT and n=10 KO mice. Student *t*-test. (*P<0.05, n.s.=non significant). SMs are significantly increased in KO CB and CX. (**F**) Dotplot depicting FC (proportional to dot size) and p-value (in grey intensity scale) of SM species detected in cerebellar samples for all analyzed tissues. Red dots represent significantly increased lipids and blue dots significantly decreased. (**F**) Bar graphs showing the average ±S.E.M of the sum of all AcCa specie concentrations per tissue in n=8 WT and n=10 KO mice. Student *t*-test. (**P<0.001, n.s.=non significant). AcCas are significantly increased only in KO CB. (**G**) Dotplot depicting FC (proportional to dot size) and p-value (in grey intensity scale) of SM species detected in cerebellar samples for all analyzed tissues. Red dots represent significantly increased lipids and blue dots significantly decreased. PL, plasma: LV, liver; CX, cortex; CB, cerebellum; Fc, fold change; AcCa, acylcarnitine; Cer, ceramide; ChE, cholesterol ester; CL, Cardiolipin; DG, diradylglycerolipid; Hex1Cer, hexosyl-1-ceramides; LPC, lysophosphatidylcholine; LPE, lysophosphatidylethanolamine; LPS, Lipopolysaccharide; LSM, lysosphingomyelin; MG, Monoradylglycerolipid; PA, phosphatidic acid; PCh, phosphatidylcholine; PE, phosphatidylethanolamine; PG, phosphatidylglycerol; PI, phosphatidylinositol; PS, phosphatidylserine; SM, sphingomyelin; SPH, sphingosine; TG, triacylglycerolipids.

Additionally, results showed that the cerebral cortex and cerebellum are the tissues with the largest amount of altered lipid species upon SNX14 depletion (Fig 5A). Using p-value < 0.05 as a cutoff, we identified 58 and 36 altered lipid species in cerebral cortices and cerebella respectively. Furthermore, only cerebellar samples clustered by genotype in principal component analysis (Fig 5B), suggesting SNX14 has a larger impact on lipid homeostasis in cerebella than in other tissues we analyzed.

Among the 58 altered lipids in cerebral cortices, 54 had lower concentrations in KOs, and 40 belong to the phosphatidylethanolamine (PE) class (Fig 5A, C, D). PEs provide fluidity and curvatures to membranes which may facilitate vesicular budding and membrane fusion essential for neuronal synaptic vesicle formation (*36*). Thus, changes in PE species may alter cerebral cortex-dependent behaviors and executive functions in SNX14 deficiency. The remaining 4 lipid species had higher concentrations in *Snx14* KO than in WT and all were SMs (Fig 5A, E, F). Similar to cerebral cortices, *Snx14* KO cerebella exhibited increased levels of SM species and decreased PE-s (Fig 5A, D, F). In addition, *Snx14* KO cerebella were distinguishable from the cortex, liver and plasma by displaying increased levels of several acylcarnitine (AcCa) species (Fig 5A, G, H). Specifically, 6 out of 16 increased lipids in *Snx14* KO cerebella, were AcCa-s. This accounted for the majority of AcCa-s detected in cerebella (6 out of 8) and resulted in an overall increase of total AcCa concentration in *Snx14* KO cerebella.

Taken together, the lipidomic analysis shows tissue specific lipid metabolite defects including cerebellar specific AcCa accumulation that is associated with the selective cerebellar neurodegeneration characteristic of SNX14 deficiency.

### SNX14 deletion impairs lipid storage *in vivo*

Under conditions of high energy demand or nutrient deprivation, AcCa-s carry fatty acids (FA) into the mitochondria for their breakdown and energy production via beta oxidation. However, elevated concentrations of AcCa can disrupt mitochondrial function and become cytotoxic. In this regard, LDs play an essential role by storing excessive FAs and preventing AcCa induced toxicity (*37*). According to this evidence, the increase of AcCa levels in *Snx14* KO cerebella could be a consequence of defects in LD biogenesis. In line with this idea, SNX14 interacts with LDs and its deficiency leads to impaired LDs in cell cultures (*22*). Thus, we investigated whether SNX14 deletion alters LD biogenesis in the cerebellum. To this end, we stained *Snx14* KO and WT mice cerebella with Bodipy 493/503 (BD493), a fluorescent dye that stains neutral lipids typically stored in LDs. As a control of LD staining by BD493 we used liver, which is a LD rich tissue specialized in FA storage in the form of TGs. As expected, we detected abundant BD493 positive LDs in WT liver sections. Furthermore, results revealed a prominent reduction of LD amounts in *Snx14* KO liver (Fig 6A). This data is consistent with the reduction of TG levels in *Snx14* KO liver we detected by lipidomics (Fig 5A and FigS5B) and suggest that SNX14 is necessary for LD biogenesis *in vivo*, at least in the liver.

**Fig6.**
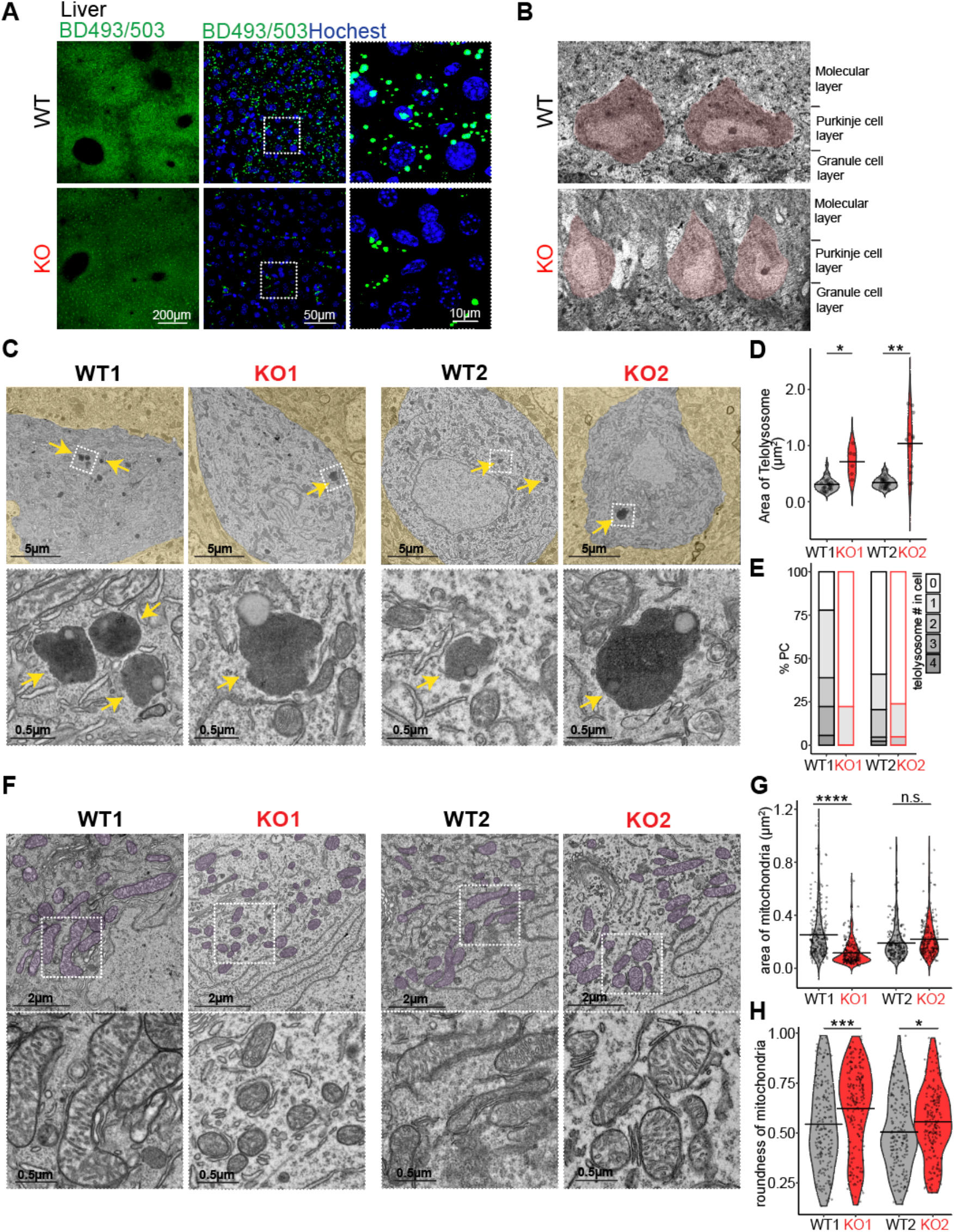
Lipid storage organelles are affected in SNX14 deficient tissue. **(A)** Representative BODIPY 493/503 (BD493) labeling shows less lipid droplets in 1-2 month old KO mice liver sections. **(B)** Representative TEM image of PC layer in WT and KO mice cerebella. **(C)** Ultrastructure analysis shows less and larger telolysosomes (yello arrow) in KO PC compared to WT. **(D)** Violine plot depicting area of telolysosomes from 15 WT1 PCs (27 telolysosomes), 6 KO1 PC (6 telolysosomes), 17 WT2 PC (30 telolysoosme), and 10 KO2 PC (12 telolysosomes). Brown-Forsythe and Welch ANOVA tests followed by Dunnett’s T3 multiple comparisons test (*P<0.05, **P<0.01). **(E)** Bar graphs showing percentage of PCs with indicated number of telolysosomes. Quantified number of PCs and telolysosomes is same as in (D). **(F)** Ultrastructure analysis shows mitochondria with intact appearance in WT and KO mice cerebellar PCs. **(G)** Violine plot depicting area of PC mitochondria from 10 PCs per animal shows significantly smaller mitochondria in one of the *Snx14* KO mice. Number of mitochondria quantified, 200 from WT1, 200 from WT2, 200 from KO1, 200 from KO2. Ordinary one-way ANOVA with Sidak’s multiple comparisons (****P<0.0001, n.s=non significant). **(H)** Violin plot depicting roundness of mitochondria reveal that KO mouse PCs have more round mitochondria, which may be indicative of mitochondrial dysfunction. The roundness of the mitochondria is defined in a scale from 0 to 1. Quantified mitochondria numbers are same as in (G). Ordinary one-way ANOVA with Sidak’s multiple comparisons. (*P<0.05, ***P<0.001).

We next focused our attention on the cerebellum. Here, BD493 staining showed few, if any, structures resembling LDs, even in WT cerebellar neurons (Fig S5C). Thus, to further explore the possibility that SNX14 deletion affects LD biogenesis in PCs, we proceeded to analyze cerebellar sections by electron microscopy (EM) (Fig 6B-H). Again, EM studies failed to identify LDs in the cerebellum. Although unconclusive, these results are not surprising if we consider that LDs are hard to detect in intact brain neurons likely due to the high turnover of FAs into membrane phospholipids (*38, 39*). Nonetheless, EM results revealed differences in lipid storage organelles between *Snx14* KO and WT PCs, including reduced number, but larger telolysosomes (Fig 6C-E). These results suggests that SNX14 may have a specialized function regulating lipid storage through the lysosomal compartment in PCs.

Finally, to determine if cerebellar AcCa increase could result in mitochondrial disfunction as previously shown (*37*), we turned our attention to mitochondrial structure. Results showed mostly intact mitochondria in *Snx14* KO PCs with slightly more round shape and smaller size in one out of the two mice analyzed. These findings are indicative of early-stage mitochondrial damage (*40*) and suggest AcCa increase may lead to PC degeneration through a mechanisms involving mitochondrial lipotoxicity.

Overall, our work shows a tissue and cell type specific sensitivity to SNX14 deficiency with a cerebellar increase of AcCa levels as the likely cause of selective PC neurodegeneration opening the windows for therapeutic alternatives through lipid metabolism regulation.

## Discussion

SNX14 deficiency causes a childhood onset cerebellar degeneration syndrome clinically defined as SCAR20 and characterized by cerebellar ataxia and intellectual disability. Previous work identified lysosome and autophagy specific defects in cultured patient neural progenitor like cells (*16, 17*) and recent evidence implicate SNX14 in lipid droplet biogenesis, fatty acid desaturation and intracellular lipid transport (*19, 22–24*). However, most of these studies were performed in cultured cells with unclear relevance for SCAR20 pathology. To overcome this limitation and study pathogenic mechanisms that selectively affect the cerebellum, we generated a *Snx14* KO mouse that closely recapitulates SCAR20 at genetic and phenotypic level. Our study shows that SNX14 deficiency leads to generalized lipid metabolite and storage defects *in vivo*. Moreover, we identify a unique profile of altered lipid species in pre-degenerating cerebella that links AcCa accumulation with selective cerebellar degeneration. Our work suggests that the role of SNX14 in lipid metabolism and storage is conserved in the nervous system and contributes to expand the emerging group of neurodegenerative disorders associated with lipid homeostasis defects.

Similar to recently reported SNX14 deficient mice(*25, 41*), the homozygous 1bp deletion in our *Snx14* KO mice causes loss of full length SNX14 protein and low RNA counts across all coding exons. However, unlike previous models that showed fully penetrant embryonic lethality (*25, 41*), ~a third of our *Snx14* KO mice develop and survive to adulthood with a phenotype that resembles SCAR20. This finding suggests that SNX14 deficiency in humans may also interrupt embryonic development, and cause SCAR20 only when embryonic lethality is circumvented. Although we still do not know what factors determine the developmental success or failure in SNX14 deficiency, there is a striking difference in the genetic architecture of *Snx14* mutations between organisms that show full and partial embryonic lethality. For instance, SNX14 deficient mice that completely fail to develop carry deletions of at least one full exon, while SCAR20 patients and animal models including our *Snx14* KO mice and previously reported dogs (*42*) and zebrafish (*25*), carry truncating point mutations or small indels. This observation has interesting implications for the generation of animal models of human disorders and for the pathogenic prediction of truncating genetic mutations that warrant further investigation.

Another factor that can influence the outcome of SNX14 deficient embryos is the environment and particularly the diet lipid composition. In line with this idea, SNX14 deficient cells are more vulnerable than control cells to saturated fatty acids (*25*) and treatment with valproic acid, which is a branched short-chain fatty acid, partially rescued PC degeneration in a conditional mouse model (*41*). Furthermore, previous studies have shown that maternal diet lipid composition can modulate brain lipidome either embryonically by maternal feeding, or in adult mice (*35*). Altogether, these data open a window to alter the course of SCAR20 through therapeutic diets. In this regard, further elucidating mechanisms that preserve lipid homeostasis in neurons, and particularly in the vulnerable PCs is of crucial relevance.

Although complex lipid homeostasis is a common pathway associated with many PC neurodegenerative disorders (*8*), very little is known about mechanisms that preserve lipid homeostasis in PCs. As neutral lipid storage sites, lipid droplets (LDs) play an important role in cellular lipid homeostasis. LDs form in response to excess nutrients or lipids, but also under cellular stress and starving conditions to store autophagy derived fatty acids and protect cells from excess fatty acid induced damage (*37, 43*). Furthermore, emerging studies show that lipid droplets play an important role in neuronal differentiation (*44*) and protecting neurons from activity-induced toxic fatty acids (*45*). In our study we failed to detect lipid droplets in cerebellar PCs despite the evident detection in the liver of WT mice and in *Snx14* KO mice at less abundance. Instead, our work revealed the presence of large but less abundant lysosome derived lipid storage organelles (i.e. telolysosomes) in *Snx14* KO mice, which suggests that PCs may have alternative lipid storage mechanisms that involve the lysosomal compartment. Notably, SNX14 has also been involved in lysosome function regulation (*16*) and recent structural predictions suggest a role in inter-organelle lipid transport (*24*) that may be important for PC specific lipid storage and homeostasis. Furthermore, recent unbiased CRISPR-based screening identified mutation of SNX14 or its paralog SNX13 as suppressors of lysosome cholesterol accumulation when the cholesterol exporter NPC1 is pharmacologically inhibited (*21*). As NPC1 mutation itself causes the neurological disease Niemann Pick Type C, these collectively underscores potential functional links between SNX14 with inter-organelle lipid metabolism and trafficking pathways.

Overall, our work highlights the relevance of lipid homeostasis for neurodegenerative disorders and disentangles a mechanism for increased susceptibility of the cerebellum to the expanding class of disorders caused by disrupted lipid metabolism pathways. Furthermore, our study provides a mouse model and molecular targets for future therapeutic studies.

## Materials and Methods

### Animals

#### General

All animal procedures were performed according to NIH guidelines and approved by the Institutional Animal Care and Use Committee (IACUC) at Children’s Hospital of Philadelphia and University of California San Diego.

#### Generation of mouse model

*Snx14* KO mice were generated by CRISPR/Cas9 editing at University of California, San Diego Transgenic Mouse Core by pronuclear injection of 5ng/ul Cas9 mRNA and 2.5ng/ul sgRNA (5’-GTAAACACGTTCTCCAAC-3’) in 1 cell stage fertilized embryos obtained from superovulated C57BL/6J females mated with C57BL/6J males. Pups were genotyped by PCR with forward 5’-cctttctgttactcagcaataacttg-3’, reverse 5’-tgaatttggaattgcgtgtg-3’ primers, followed by Sanger sequencing and those carrying *Snx14* indel alleles selected for backcross with WT C57BL/6J mice for 3-6 generations (to filter out potential off targets) and further expanded as an experimental model. Only the Snx14 c.1432delG carriers generated homozygous pups.

#### Animal maintenance and husbandry

Mice were group-housed at a maximum 5 animals per cage with a 12-h light/dark cycle at constant temperature with ad libitum access to food and water and maintained in C57BL/6J background (C57BL/6J, Jackson Laboratories Stock No: #000664). To obtain homozygous *Snx14* KO mice and WT littermates, heterozygous male and female mice were mated. Genotyping was performed by qPCR following Transnetyx Inc. (Cordova, TN) custom protocol with DNA obtained from tail clips.

### Behavior analysis

#### Experimental design

Behavior analysis was performed with three cohorts of WT and *Snx14* KO littermates starting at 8 months of age. Each cohort contained mixed genotype and sex of animals. Behavior tests were performed in the following order: accelerating Rotarod, Catwalk, Metz Ladder and Social choice/recall. Investigators were blinded during scoring of behavioral assessments. Whenever possible, offline analysis by computer software was utilized to enhance rigor.

#### Accelerating Rotarod

On day 1, mice were habituated to the stationary Rotarod for 2 minutes. This was immediately followed by a trial where rotation was programmed to rise from 4-40rpm in 300 seconds. After a 30 minute intertrial interval (ITI), a second trial was performed, followed by another ITI and third trials. Three additional trials were performed on the next 2 consecutive days, for a total of 9 trials. A trial was terminated when a mouse fell, made one complete revolution while hanging onto the rod, or after 300s. Latency to fail (time stayed until falling or riding the rod for a single revolution) was determined. Learning rate was calculated as followed: learning rate = (Trial 9 latency to fall – Trial 1 latency to fall)/8, 8 is the number of inter-trial intervals in this study.

#### Catwalk gait analysis

In the Catwalk gait analysis assay, mice were placed on a meter-long illuminated glass plate walkway in a dark room. A high-speed video camera below the plate recorded the paw prints, as the mice traversed a 20cm section of the alley. The paw print footage was analyzed by CatWalk XT program (Noldus, Leesburg, VA).

#### Metz ladder rung waking test

The Metz procedure used a 1 meter long horizontal ladder, which was about 1cm wider than the mice. The Plexiglas walls were drilled with 3mm holes to accept the metal rungs. The gaps between the rungs were randomly spaced 1-5 cm apart so that the mice had to adjust the projection of the landing of each paw. Mice were trained to run the ladder with all rungs in place, 1cm apart before the test trials began. In the test, each mouse was placed at the beginning of the ladder. Five trials were performed on consecutive days and videotaped. The pattern of the rungs was changed after each trial to prevent animals from adapting. Trials were recorded by a high definition digital camera. Foot slip(s) of each trial was quantified later by an investigator blinded to group designation with video.

#### Social choice and recall test

Mice were tested for social preference and recall as described previously (*46*). The testing apparatus was a rectangular Plexiglas three chamber arena (60 cm (L) × 40 cm (W) × 20 cm (H)). The chamber was continuous with areas at the ends designated for the placement of vented cylinders to hold the cues. The social cues were juvenile, sex-matched C57BL/6J mice. The inanimate cues were smooth rocks that approximate the size of the social cues. The procedure consisted of a habituation phase whereby the experimental mouse was placed into the center chamber with empty cylinders in the side chambers for 10 minutes. After habituation, the choice phase immediately began. The cylinders were loaded with either a social cue (young mouse, M1) or inanimate cue. The experimental mouse was allowed to explore the cues for 10 minutes. Immediately after the choice phase, the recall phase was performed. The now familiar social cue, M1 remained in a cylinder while a novel mouse, M2 was loaded into the cylinder that previously held the inanimate cue. The experimental mouse was allowed to freely explore the 2 social cues for 10 min. The bouts and duration of explorations (nose <1 cm proximity) with the cylinderswas determined with ANYmaze software (Stoelting Co. Wood Dale Il.).

### Histology

#### Immunofluorescence staining

Mice were anesthetized with isoflurane (Terrell) and perfused trans-cardially with 20ml PBS and 20ml 4% paraformaldehyde (Electron Microscopy Sciences). Brains dissected out from scalp were post-fixed in 4% paraformaldehyde for 18h in RT and washed 3 x 10 mins in PBS. Brains were sliced into 50um sections using vibratome (Leica).

On the day of staining, slides were washed with PBS, permeabilized and blocked with PBS+0.3% Triton X-100 (PBST) and 5% goat serum (G9023, Sigma-Aldrich). Slides were then incubated with primary antibodies in 5% goat serum in PBST at 4°C on the shaker overnight. Primary antibodies used include Calbindin-1 (1:500, C9848, Sigma-Aldrich), GFAP (1:500, Z033429-2, Agilent, Santa Clara, CA), and IBA1 (1:500, 019-19741, Wako, Osaka, Japan), NeuN (Sigma-Aldrich, MAB377). Next day, slides were washed with PBST 3 x 10 mins and incubated with Alexa Fluor-conjugated secondary antibodies at 1:500 in 2% normal goat serum in PBST for 2h at room temperature (RT). Secondary antibodies used include Alexa Fluor 488-labeled goat anti-mouse IgGs (A-11001, Invitrogen), Alexa Fluor 555-labeled goat anti-mouse IgGs (A-21422, Invitrogen), Alexa Fluor 488-labeled goat anti-rabbit IgGs (A-11008, Invitrogen), and Alexa Fluor 555-labeled goat anti-rabbit IgGs (A-21428, Invitrogen). Slides were washed in PBST 3 x 10 mins, incubated with 300nM DAPI (D3571, Invitrogen) for 10min at RT and mounted in microscope slides with ProLong Gold antifade (P36930, Invitrogen) or Mowiol (sigma #81381) covered with coverslip. Immunostainings were imaged with a Leica TCS SP8 X confocal microscope and images processed and quantified with ImageJ (NIH).

#### BODIPY staining

Mice were perfused as above, and brain and liver dissected out and post-fixed in 4% paraformaldehyde for 18h in RT following three washes with PBS. Tissue was sliced into 50μm sections using vibratome (Leica), rinsed in PBS and incubated with 2μM BODIPY 493/503 (Invitrogen D3922) for 30 min at RT with gentle rocking. Then, the sections were rinsed in PBS 3 x 10 mins and mounted on microscope slides with Mowiol (sigma #81381) covered with coverslips.

#### Transmission Electron microscopy

Mice were perfused with 20ml of PBS, followed by 20ml 2% paraformaldehyde and 2% glutaraldehyde in sodium cacodylate buffer. Then, cerebella were dissected out, trimmed to 1mm thickness, and processed for transmission electron microscopy in Delaware University Delaware Biotechnology Institute. Briefly, tissues were washed 3 x 15 min in 0.1M sodium cacodylate buffer pH 7.4 and post-fixed for 2 h with freshly prepared 1% osmium tetroxide and 1.5% potassium ferrocyanide in 0.1M sodium cacodylate buffer pH 7.4. The tissue was washed 4 x 15 min with water and *en bloc* stained overnight with 1% uranyl acetate (aq) and washed again with water. The samples were dehydrated through an ascending acetone series (25%, 50%, 75%, 95%, anhydrous 100%, anhydrous 100%), 15 min each step, and then infiltrated with Spurr low viscosity resin (25% resin in acetone, 50% resin in acetone, 75% resin in acetone) for 1 h each step. Following several changes in 100% resin, the tissue infiltrated overnight on a rotator, and the next day, samples were embedded in aluminum weigh dishes and polymerized at 60°C overnight. Ultrathin sections were cut using a Leica UC7 ultramicrotome, and sections were placed onto 100 mesh formvar/carbon coated copper grids. Sections were post-stained with 2% uranyl acetate in 50% methanol and Reynolds’ lead citrate. Sections were examined on a Zeiss Libra 120 transmission electron microscope operating at 120kV, and images were acquired with a Gatan Ultrascan 1000 CCD. Quantification of area and numbers was done by ImageJ (NIH) and graphs generated with R 4.2.

### Biochemical studies

#### Western blot

Mouse tissue was dissected, fast-froze, and stored in −80°C until use if needed. On the experiment day, tissue was homogenized in RIPA buffer (#9806, Cell Signaling) supplemented with protease inhibitor cocktail (P8340, Sigma-Aldrich) with microtube homogenizers and incubated for 15 minutes at 4C. After centrifugation at 13,200 rpm, supernatant containing protein extract was collected, mixed with 1xLDS loading buffer (B0007, Invitrogen) supplemented with 200mM DTT (BP172-5, Fisher Scientific) and loaded on a 4-15% Mini-Protean TGX Precast Protein Gel. Proteins were transferred onto PVDF membranes in Mini Gel Tank at 80V for 180min. Membranes were blocked with 5% milk-TBST for 1h at room temperature (RT) then probed with primary antibodies diluted in 5% milk-TBST solution overnight at 4°C. Primary antibodies used include SNX14 (1:1000, HPA017639, Sigma-Aldrich, St. Louis, MO) and Beta Actin (1:2000, A00702, GenScript, Piscataway, NJ). Membranes were then washed and probed with horseradish-peroxidase-conjugated donkey anti-mouse or anti-rabbit IgG (H+L) cross-adsorbed secondary antibody (SA1-100, Invitrogen; 31438, Invitrogen) for 1h at RT. Membranes were developed using SuperSignal™ west dura extended duration substrate (34076, Invitrogen) and exposed on film. Exposed films were scanned, and protein bands were quantified using ImageJ Software (NIH).

#### RNA-seq

1 month old or 1 year old mice were euthanized and tissue was dissected on ice, fast frozen, and stored in −80°C until RNA extraction. On the day of RNA extraction, 50-100mg tissue from each sample was lysed in 1ml TRIzol (15596026, Invitrogen). Addition of 0.2ml chloroform (288306, Sigma-Aldreich) followed by centrifugation was performed to separate the solution into an aqueous phase and an organic phase. Aqueous phase was collected and Isopropyl alcohol was added to precipitate RNA. After centrifugation pellets were washed with 75% ethanol (111000200CSPP, Pharmco) and resuspended in RNAse free water. Strand-specific mRNA-seq libraries for the Illumina platform were generated and sequenced at GENEWIZ or Novogene following the manufacturer’s protocol with sample specific barcodes for pooled sequencing. After sequencing in Illumina HiSeq or Novoseq platform with 2×150 PE configuration at an average of 15 million reads per sample, sequenced reads were trimmed to remove possible adapter sequences and poor quality nucleotides and trimmed reads mapped to the Mus musculus GRCm38 reference genome using Spliced Transcripts Alignment to a Reference (STAR) software. Unique hit counts were then extracted using featureCount from the Subread Package v.1.5.2. and differential expression analysis performed in DESeq2 *(47)*. Volcano plots were generated with R package *ggplot2.* Gene ontology enrichment was calculated with datasets from Molecular Signature Database (MsigDB) (v7.1) under the tag *GO biological process (BP)*. Circular plot generated with R package GOplot (*48*).

#### UPLC-HRMS whole lipidome analysis

##### Sample preparation

1-2 mo mice were euthanized and, after heart blood collection, cortex, cerebellum and liver were dissected and snap-frozen in liquid nitrogen. Blood, collected in lavender top tubes (RAM Scientific 07 6011), was centrifuged at 2000g for 15min at 4°C and the supernatant (EDTA plasma) collected and snap-frozen in liquid nitrogen. Snap-frozen plasma and tissue were stored at −80 until lipid extraction. For lipid extraction, plasma samples were prepared as previously reported (*49*) and ~10mg of frozen tissue fragments were weighted and chopped with dry-ice-chilled blades on a chilled tile. The tissue was added to low retention Eppendorf tube filled with 0.6 mL 80% methanol (MeOH) and 10 μL on internal standard mix (SPLASH® LIPIDOMIX #330707 from Avanti Polar Lipids, Alabaster, AL) and kept in dry ice. Samples were pulse sonicated in ice for 30x 0.5 second and incubated for additional 20 min in ice for metabolite extraction. Each tube was then vortexed 3x 30 seconds and tissue homogenates transferred to a 10 mL glass Pyrex tube with screw cap. The Eppendorf tubes were rinsed with 0.5 mL methanol and added to corresponding glass Pyrex tube. Then, 5 mL methyl tert-butyl ether (MTBE) was added to each tube and vigorously shacken for 30 minutes, followed by the addition of 1.2 mL water and 30 second vortex. Samples were centrifuged for 10 min at 1000g to created two phases. The top clear phase was collected to a clean glass Pyrex tube and dried down under nitrogen. For the analysis, dried samples were resuspended in 100 μL MTBE/MeOH=1/3 (v/v), spun down at 10,000g for 10 min at 4°C. The top 50 μL were transferred to a HPLC vial and 2ul were injected for LC-MS analysis.

##### Liquid chromatography high resolution -mass spectrometry (LC-HRMS) for lipids

Separations were conducted on an Ultimate 3000 (Thermo Fisher Scientific) using an Ascentis Express C18, 2.1 × 150 mm 2.7μm column (Sigma-Aldrich, St. Louis, MO). Briefly, the flow-rate was 0.4 m/min, solvent A was water:acetonitrile (4:6 v/v) with 0.1% formic acid and 10 mM ammonium formate and solvent B was acetonitrile:isopropanol (1:9 v/v) with 0.1% formic acid and 10 mM ammonium formate. The gradient was as follows: 10 % B at 0 min, 10 % B at 1 min, 40 % B at 4 min, 75 % B at 12 min, 99 % B at 21 min, 99 % B at 24 min, 10 % B at 24.5 min, 10 % at 30 min. Separations were performed at 55 °C.

For the HRMS analysis, a recently calibrated QE Exactive-HF mass spectrometer (Thermo Fisher Scientific) was used in positive ion mode with an HESI source. The operating conditions were: spray voltage at 3.5 kV; capillary temperature at 285°C; auxiliary temperature 370°C; tube lens 45. Nitrogen was used as the sheath gas at 45 units, the auxiliary gas at 10 units and sweep gas was 2 units. Same MS conditions were used in negative ionization mode, but with a spray voltage at 3.2 kV. Control extraction blanks were made in the same way using just the solvents instead of the tissue homogenate. The control blanks were used for the exclusion list with a threshold feature intensity set at 1e10^5. Untargeted analysis and targeted peak integration was conducted using LipidsSearch 4.2 (Thermo Fisher Scientific) as described by Wang et al (*50*).

An external mass calibration was performed using the standard calibration mixture approximately every three days. All samples were analyzed in a randomized order in full scan MS that alternated with MS2 of top 20, with HCD scans at 30, 45 or 60 eV. Full scan resolution was set to 120,000 in the scan range between m/z 250–1800. The pool sample was run every 15 samples. Lipids quantification was done from the full scan data. The areas were normalized based on the amount of the internal standard added for each class. All amounts were then normalized to the original tissue weight.

### Statistics

Statistical analyses were performed using GraphPad Prism 8 (GraphPad Software, Inc., La Jolla, CA). When possible, data was analyzed blind to the genotype. Sample size for each experiment was determined based on similar studies. To compare the means of groups where normal distribution and similar variance between groups was confirmed, Student’s t test (for two samples), one-way ANOVA (for more than two samples) or two-way ANOVA followed by Tukey’s post hoc test (for multiple variables) was be used. Pval<0.05 was used as cutoff for significance and data represented as bar graphs, curves or scattered dot plots. Outliers were removed in two behavioral studies using the using ROUT method with Q=1%, p<0.0002.

## Acknowledgments

We thank Shannon Modla at Delaware University Biotechnology Institute electron microscopy core for processing and imaging tissue for transmission electron microscopy and the UCSD transgenic core for the pronuclear injections to generate *Snx14* KO mice. Behavior procedures were performed with assistance from The Neurobehavior Testing Core at UPenn, ITMAT and IDDRC at CHOP/Penn.

## Author contributions

Study conceptualization and design: Y.Z. and N.A. Validation and maintenance of mouse colony: Y.Z. and N.A. Behavioral study design, execution, and data collection: B.C. and T.O. Behavioral data analysis: B.C., T.O., Y.Z. and H.T. Histology and TEM studies: Y.Z., V.S., M.F. D.Y. RNA extraction and RNAseq analysis: Y.Z. Lipidomic analysis: Y.Z., P.X., and C.M. Data interpretation: Y.Z., M.H. and N.A. Supervision and project administration: N.A. Manuscript preparation: Y.Z. and N.A. Manuscript edit and review: All authors

## Data and materials availability

RNAseq data was deposited in GEO under the GSE215834 reference. All the other data are available in the main text or the supplementary materials.

## Supplementary Materials

**Fig. S1:**
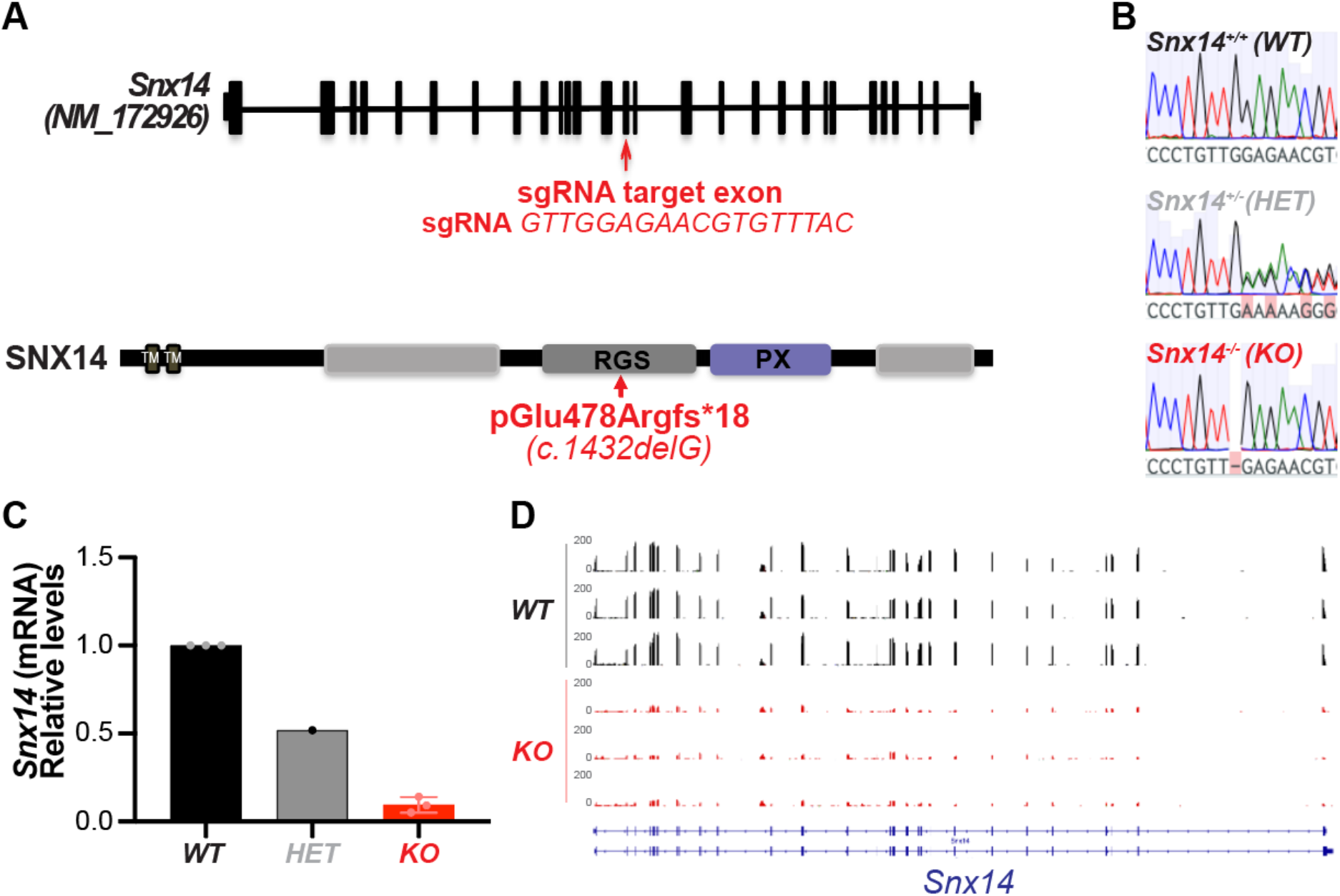
1bp deletion in exon 14 of *Snx14* causes loss of SNX14 expression. **(A)** Diagram showing the sgRNA sequence used to target exon14 with CRISPR/Cas9 (top) and the mutation carried by our *Snx14* KO mice. **(B)** Representative Sanger sequencing chromatograms of *Snx14*^+/+^ (WT), *Snx14*^+/−^ (HET), and *Snx14*^−/-^ (KO) mouse PCR product, showing the c.1432delG homozygous deletion in a *Snx14* KO mouse. **(C)** RT-qPCR of *Snx14* in WT, HET, and KO mouse tissues, show reduction of *Snx14* mRNA levels in KO mice. **(D)** Integrative Genomics Viewer (IGV) screenshot of RNAseq results showing less abundant read counts across all the *Snx14* exons in *Snx14* KO than in WT mice. Bottom blue diagrams show two alternative splice forms of *Snx14*. Thin lines represent introns and thick bars exons.

**Fig. S2.**
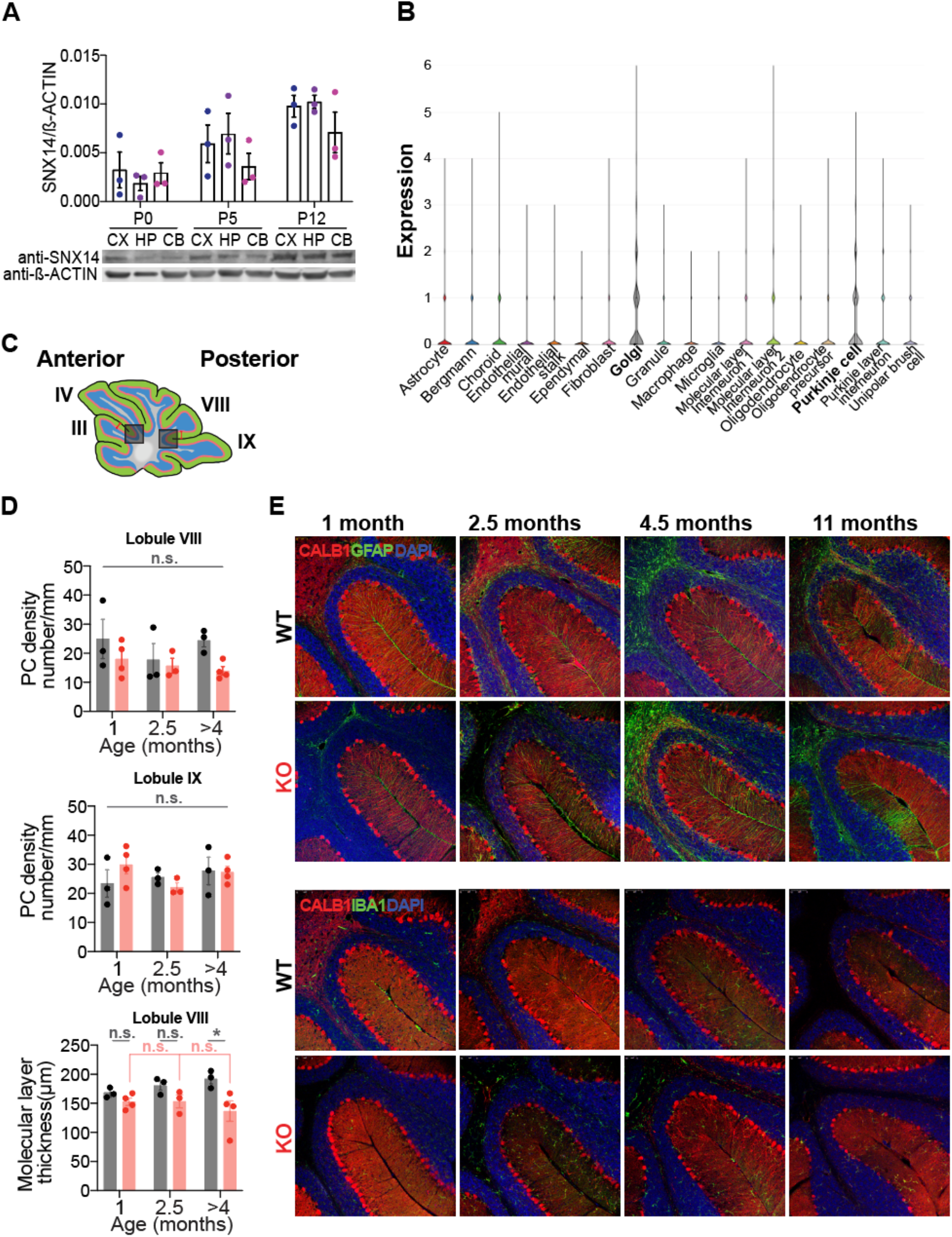
SNX14 is widely expressed and its deficiency causes to late onset degeneration of posterior cerebella. **(A)** Western blot results showing the expression level of SNX14 in various brain regions in early postnatal ages (bottom). Graph shows relative SNX14 expression quantified by WB band densitometry of n=3 mice for each time point. CX, cortex; HP, hippocampus; CB, cerebellum. βActin is shown as loading control. (**B**) Violin plots showing distribution of cerebellar cells based on their *Snx14* expression levels. Data was extracted from cerebellar single nuclei (sn) RNAseq *(30)* deposited in Broad Institute Single Cell Portal. (**C**) Diagram showing cerebellar regions depicted in Fig 3D, G, H and Fig S2D, E immunofluorescence images. (**D**) PC density quantified from Lobule VIII and IX, and molecular layer thickness quantified from Lobule VIII. 1mon, WT n=3, KO n=4; 2.5mon, WT n=3, KO n=3; >4mon, WT n=3, KO n=4. Two-way ANOVA followed by Sidak test. (*P<0.05, n.s.=non significant). (**E**) Immunofluorescent staining labeling Purkinje cells in red with anti-Calbindin antibody and astrocytes in green with anti-GFAP antibody (top) and microglia in green with anti-IBA1 antibody (bottom) at the base of Lobule VIII & IX.

**Fig. S3.**
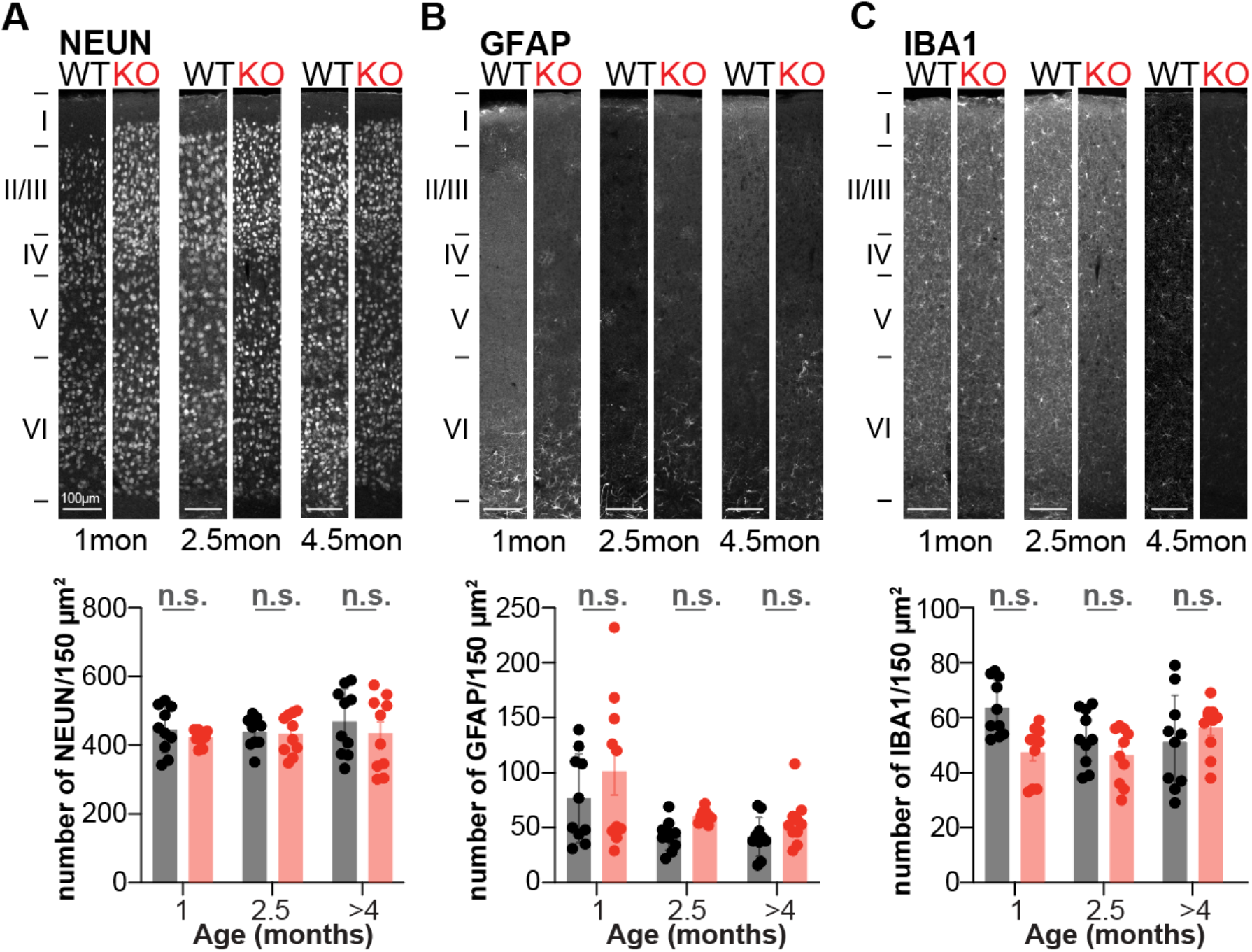
No signs of neurodegeneration in *Snx14* KO cerebral cortex. (**A-C**) Coronal sections of cerebral cortices immunostained with (A) anti-NeuN to label neurons (B) anti-GFAP to label astrocytes and (C) anti-IBA1 to label microglia. Bar graphs showing average ±S.E.M of cell numbers per 150um^2^ area in 4-5 regions of 3 mice per genotype and age. Two-way ANOVA followed by Sidak test (n.s.=non significant).

**Fig. S4.**
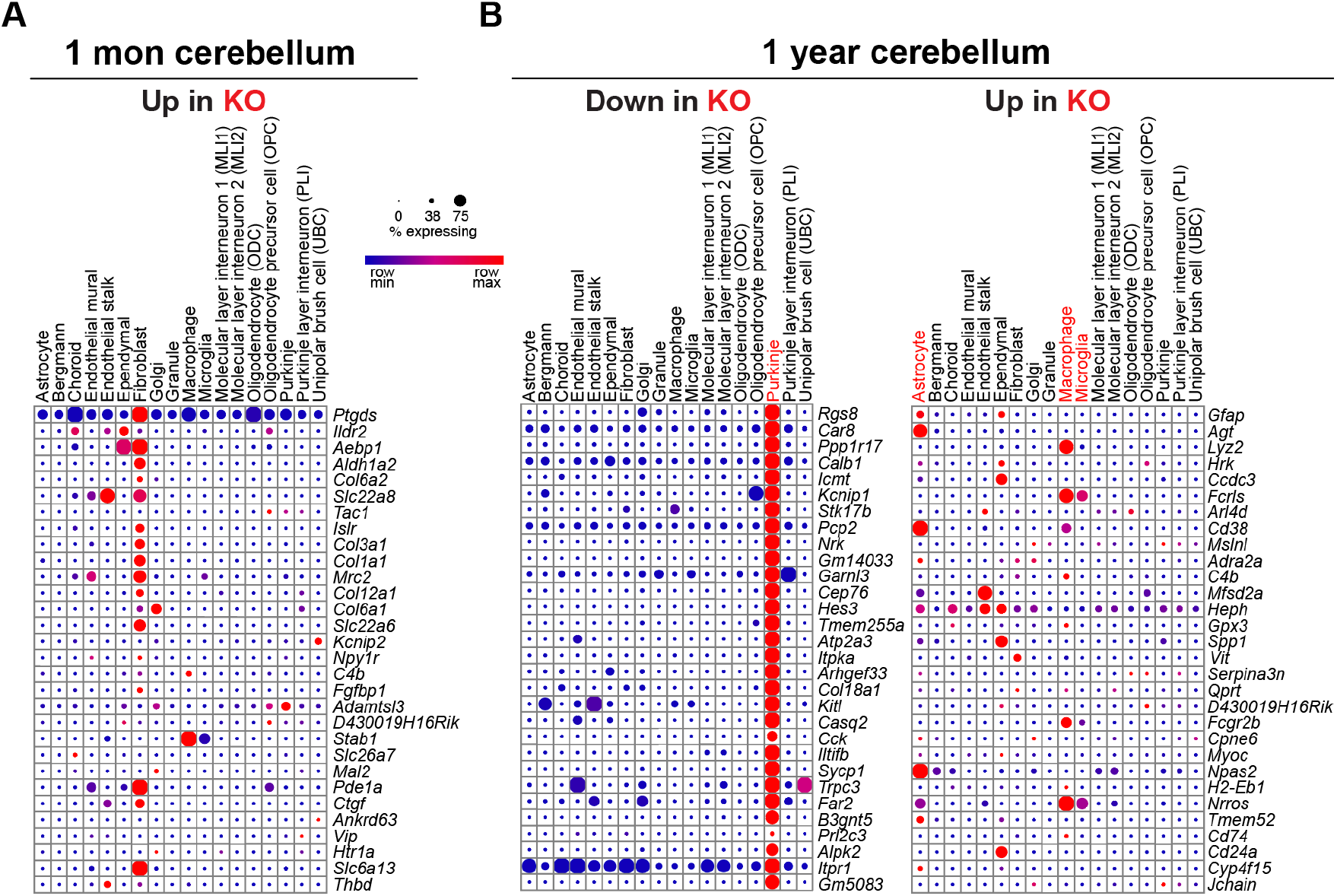
Loss of PC specific gene expression in 1-year-old *Snx14* KO cerebella. (**A**) Dot plot of cerebellar snRNAseq data *(30)* showing the average expression level (in blue to red scale) and percentage of cells (dot size) expressing the top 30 upregulated DEGs in 1 month old *Snx14* KO cerebella (from our bulk RNAseq results). (**B**) Same as in (A) for the top 30 downregulated DEGs (left), and top 30 upregulated DEGs (right) in 1-year-old *Snx14* KO cerebella. Data shows that downregulated DEGs in 1-year-old *Snx14* KO are typically expressed in PCs which indicating that there is a loss of PCs in *Snx14* KO cerebella.

**Fig. S5.**
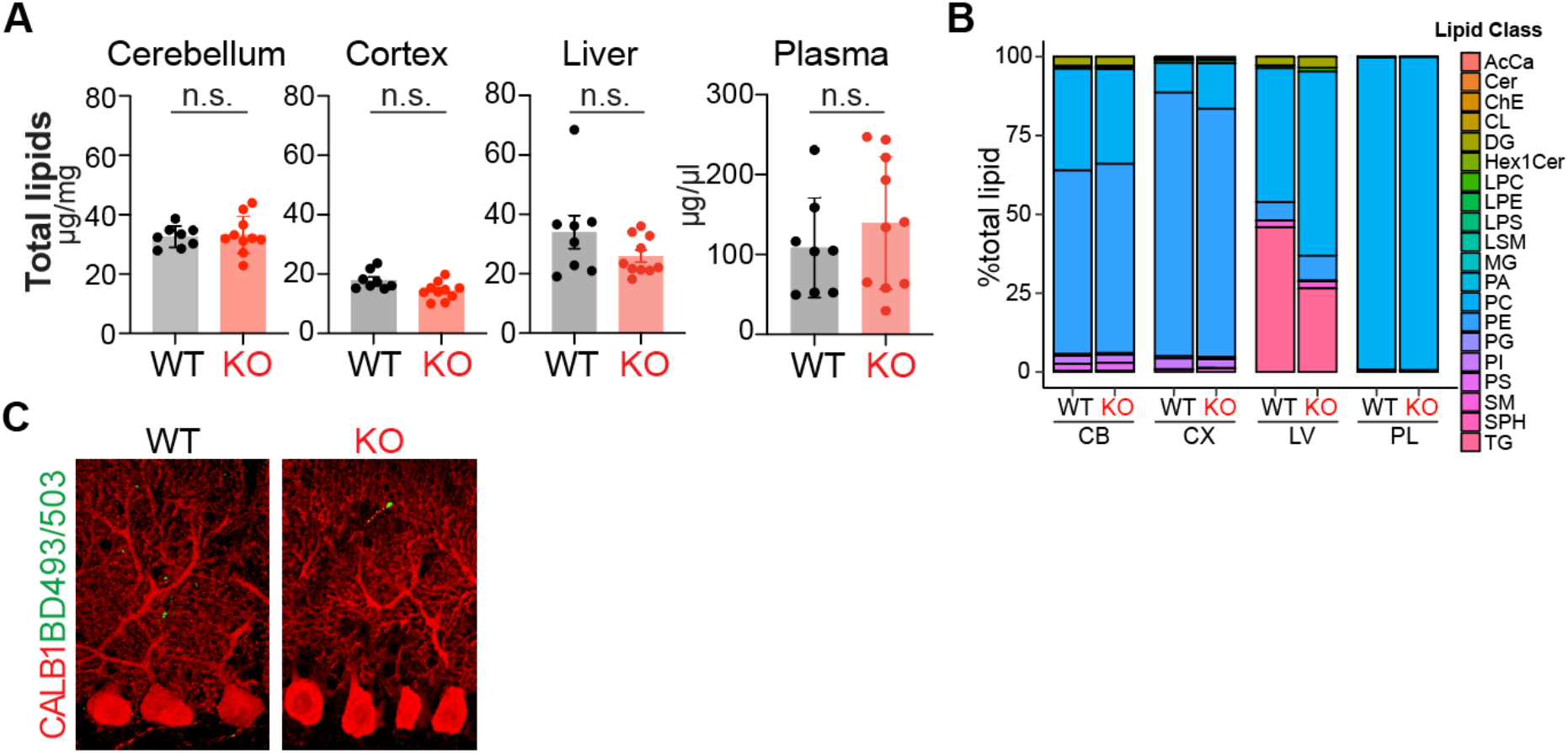
SNX14 deficiency alters lipid metabolite abundance. (**A**) Bar graphs showing average ±S.E.M of total lipid concentration in cerebellum, cortex, liver and plasma of n=8WT and n=10KO mice. Student t-test (n.s.=non significant). (**B**) Percentage of each lipid class in indicated tissue of WT and KO mice. (**C**) Representative image of BD493/503 lipid droplet staining (green) in CALB1 positive PCs (red) of WT and KO cerebellar sections. AcCa, acylcarnitine; Cer, ceramide; ChE, cholesterol ester; CL, Cardiolipin; DG, diradylglycerolipid; Hex1Cer, hexosyl-1-ceramides; LPC, lysophosphatidylcholine; LPE, lysophosphatidylethanolamine; LPS, Lipopolysaccharide; LSM, lysosphingomyelin; MG, Monoradylglycerolipid; PA, phosphatidic acid; PCh, phosphatidylcholine; PE, phosphatidylethanolamine; PG, phosphatidylglycerol; PI, phosphatidylinositol; PS, phosphatidylserine; SM, sphingomyelin; SPH, sphingosine; TG, triacylglycerolipids.

